# Altered cortical development and psychiatric symptom risk in adolescents exposed to maternal stress *in utero*

**DOI:** 10.1101/540930

**Authors:** Goldie A. McQuaid, Valerie L. Darcey, Melissa F. Avalos, Diana H. Fishbein, John W. VanMeter

## Abstract

**Objective:** Maternal exposure to stress during pregnancy is associated with increased risk for cognitive and behavioral sequelae in offspring. Animal research demonstrates exposure to stress during gestation has effects on brain structure. In humans, however, little is known about the enduring effects of *in utero* exposure to maternal stress on brain morphology. We examine whether maternal report of stressful events during pregnancy is associated with brain structure and behavior in adolescents.

**Method:** We compare gray matter morphometry of typically-developing early adolescents (11-14 years of age, mean 12.7) based on presence/absence of maternal report of stressful event(s) during pregnancy: prenatal stress (PS; n=28), no prenatal stress (NS; n=55). The Drug Use Survey Index Revised (DUSI-R) assessed adolescent risk for problematic behaviors. Exclusionary criteria included pre-term birth, low birth weight, and maternal substance use during pregnancy. Groups were equivalent for demographic (age, sex, IQ, SES, race/ethnicity), and birth measures (weight, length).

**Results:** Compared to NS peers, adolescents in the PS group exhibited increased gray matter volume in bilateral posterior parietal cortex (PPC): bilateral intraparietal sulcus, left superior parietal lobule and inferior parietal lobule. Additionally, the PS group displayed greater risk for psychiatric symptoms and family system dysfunction, as assessed via DUSI-R subscales.

**Conclusion:** This study provides evidence for the enduring neurodevelopmental effects of exposure to maternal stress while *in utero*, suggesting the dynamic alterations in brain structure that occur during the adolescent critical period may unmask the latent effects of gestational stress on brain morphology and increase subsequent risk for psychiatric symptomatology.

## INTRODUCTION

A large body of animal research has demonstrated negative effects of prenatal stress on the structure of the developing brain in offspring, including preclinical^1–3^ and non-human primate studies^4,5^. *In utero* stress exposure has adverse effects on neurodevelopment of the corpus callosum^4^, hippocampus^5–7^, amygdala^8^, rostral anterior cingulate cortex^9^, and frontal cortex^10^. Corresponding closely with the functional neuroanatomy of these regions, offspring of animals stressed during gestation exhibit deficits in attention^11^, as well as an increase in behaviors associated with anxiety^12,13^ and depression (e.g., altered circadian rhythms^14^ and learned helplessness^15^). Critically, the negative effects of prenatal stress endure throughout the lifespan of the animal^16^.

As in animals, research in humans has demonstrated that maternal exposure to stress during pregnancy is associated with adverse cognitive and behavioral outcomes in childhood, adolescence, and adulthood. These studies operationalize stress in various ways, including major life events^17^, natural disasters^18,19^, military invasion^20^, and maternal psychiatric symptoms, such as depression^21^ and anxiety^22,23^. Despite differences in methodologies and specific findings, research reveals a largely consistent picture showing maternal stress experienced during pregnancy is related to aberrant cognitive, behavioral and emotional development in offspring, and is associated with heightened risk for development of psychiatric symptoms and illnesses, including impulsivity^22^, attention deficit hyperactivity disorder ^24,25^, anxiety ^21^, depression^23^, autism spectrum disorder^17,26^, and schizophrenia^20,27,28^.

While effects of maternal stress on cognition and behavior are thoroughly documented, studies have only recently examined impacts on child neurodevelopment. These nascent studies have demonstrated effects of *in utero* stress on white matter organization^29,30^, and on gray matter in cortical^31–33^ and subcortical^34^ structures. For instance, in neonates, prenatal maternal anxiety is associated with reduced cortical thickness in frontal and parietal cortices, an effect modulated by genetic variants implicated in increased vulnerability to psychiatric disorders^35^. Effects of maternal stress during gestation have been observed in older offspring as well. Maternal reports of stress during pregnancy are associated with differences in white matter microstructural organization of the uncinate fasciculus in offspring aged 6-9 years^29^. Further, studies of 6-9 year-olds have shown that greater maternal pregnancy-specific anxiety at 19-weeks gestation is linked to reduced gray matter volume in bilateral prefrontal cortex, temporal and parietal cortices, and cerebellum^31^, and greater maternal depressive symptoms are associated with thinner cortices across the brain^32^. Finally, maternal first-trimester cortisol (a biological marker of stress) is positively correlated with right amygdalar volume in 7-year old female offspring; further amygdalar volume mediates the relationship between higher maternal cortisol and increased affective symptoms in female offspring^34^.

The trajectory of cortical maturation from childhood through adulthood identifies adolescence as a critical period in the non-linear development of frontal and parietal cortical gray matter^36–38^. It is important to note that during adolescence gray matter volume is expected to decrease as a result of cortical pruning. Cortical gray matter volume during this developmental period may be a relatively sensitive indicator of earlier neurodevelopmental influences, and previously masked deficits may be revealed by the maturational processes occurring during this time^39^. Understanding how prenatal stress exposure impacts the developmental trajectory of cortical development during the adolescent period may be particularly revealing, as many types of mental health problems emerge in adolescence^40^. Adversity early in life has been shown to set forth a “cascade” of effects in the developing brain^41–43^, including those at the molecular, cellular and network levels, which may only manifest during a later developmental period^44–46^, such as adolescence.

Although the mechanisms by which gestational stress impacts offspring human neurodevelopment and increases risk for psychiatric symptoms have yet to be fully elucidated^47–49^, both the animal and human literature demonstrate that prenatal stress affects brain development and can play a contributory etiological role in cognitive and behavioral sequelae. Identifying offspring brain structures affected by prenatal stress exposure and understanding how these structures are altered, as well as the behavioral and psychiatric outcomes of these structural alterations, may allow for identification of youth at elevated risk for poor outcomes.

The current study examined cortical gray matter volume in two groups of typically developing, early adolescents who differed with regards to whether mothers experienced at least one major stressful life event during their pregnancy. While we did not predict specific neuroanatomical regions to differ *a priori*, we hypothesized that adolescents whose mothers experienced significant stress during their pregnancy would have a neuroanatomical profile suggestive of delayed gray matter development compared to their non-prenatally stressed peers.

## METHODS

Participants were recruited as part of the Adolescent Development Study, a prospective longitudinal investigation of the neural, cognitive and behavioral precursors and consequences of substance use initiation and escalation. Demographic, cognitive, behavioral, and imaging assessments were conducted at baseline and were repeated during two follow-up visits, approximately 18- and 36-months later. The data reported here were collected during the baseline visit. Complete information on the methods of this study are described in detail elsewhere^50^. Briefly, 135 substance-naïve 11-13-year-olds and their caregivers were recruited from the Washington D.C. metropolitan region. Primary exclusionary criteria were prior substance use by the adolescent, neurodevelopmental disorder (e.g., autism spectrum disorder, active tic disorder that would interfere with imaging), prior head trauma, left-handedness, and conditions contraindicated in MRI. The Georgetown University Institutional Review Board approved the study, and caregivers and adolescents provided consent and assent prior to all data collection.

### Family/caregiver measures

#### Pre-/Perinatal questionnaire

Biological mothers were invited to complete a pre-/perinatal questionnaire collecting information about the pregnancy with, and delivery of, their enrolled adolescent. Mothers were asked to indicate whether any stressful events occurred during their pregnancy with the adolescent participant. Questions concerning stressors were adapted from the second item of a short screening scale for posttraumatic stress disorder (PTSD)^51^. Items from the scale probing PTSD symptom domains (e.g., re-experiencing of traumatic event(s), exaggerated startle response, etc.) were omitted. The following stressful events were queried: relationship conflict (e.g., divorce, break up, infidelity), death of someone close (e.g., partner, parent, another child), severe illness/harm of someone close (e.g., cancer, heart attack, severe car accident), severe financial issues (e.g., major property damage, foreclosure, sudden unemployment), involvement in an accident (e.g., car, fire, other disaster), being physically or emotionally attacked (e.g., threatened with a weapon, rape), or another terrible event that most people do not experience (with space provided for elaboration). Mothers could indicate that they had experienced no stressful event during pregnancy by selecting the option “nothing like this happened to me while I was pregnant with this child.”

Mothers also indicated their age at time of delivery, incidence of any obstetric complications, and extent of any tobacco, alcohol, or prescription or illicit drug use during the pregnancy. Finally, outcomes after pregnancy were reported for both the mother (post-partum depression(PPD)) and child (birth weight and length).

#### Family socioeconomic status index (SES)

An SES Index was calculated by averaging the mean of two standard scores (mean household income bracket before taxes and mean cumulative parental education), and re-standardizing these to obtain a z-score distribution with a 0-centered mean and a standard deviation of 1 for the sample analyzed (n=83) (method adapted from^52^).

### Adolescent measures

#### Risk measure

Adolescents completed the Drug Use Survey Index Revised (DUSI-R), a 159-item survey that measures current use of drugs and alcohol, risk for future problematic use, as well as experiences and behaviors known to precede and co-occur with substance use^53,54^. Although non-diagnostic in nature, previous research has shown the DUSI-R to be psychometrically valid for identifying adolescents at risk for adverse outcomes, including disrupted psychosocial development, psychiatric symptomatology, and antisocial behavior^55–60^. Survey questions are *yes-no* items categorized into ten domains: psychiatric symptoms, family system, behavior patterns, health status, leisure and recreation, social competence, school performance, peer relationships, substance use, and work performance. For each domain a problem density index was tabulated^59^. Higher scores suggest greater problems within a particular domain. Given limited variability and null responses (by design) in two of the subscales (work performance and substance use), the present analysis examines the other 8 DUSI subscales.

#### Intelligence and physical development

Full-scale intelligence was estimated via the Kaufman Brief Intelligence Test, 2^nd^ edition (KBIT-2)^61^. In order to determine physical growth characteristics in the adolescent groups, pubertal development and body mass index (BMI z-score) were assessed. Participants completed the Scale of Physical Development^62^, a self-report questionnaire highly correlated with Tanner stage^63^. The questionnaire evaluates the degree to which particular physical maturational changes have occurred, including changes in skin/voice, breast development and facial hair. Possible scores range from 1 (prepubertal) to 4 (postpubertal). BMI (kg/m^2^) was calculated using height measured via stadiometer (SECA 216 wall-mount mechanical measuring rod; triplicate measures within 0.5 cm, averaged) and weight measured via digital scale (Health-O-Meter Professional 394KLX). BMI norms for age and sex were used to determine z-scores and percentiles^64^. These three measures were collected during the baseline visit except in the case of one adolescent, whose IQ was assessed during the 1.5-year follow-up visit.

### MRI acquisition and analysis

#### MRI data acquisition

During the baseline visit, high-resolution structural images were acquired on a Siemens TIM Trio 3 T scanner with a 12-channel head coil using a T1-weighted magnetization prepared rapid acquisition gradient echo (MPRAGE) sequence. A total of 176 sagittal slices were collected using the following parameters: TR/TE/TI = 920/2.52/900 ms, flip angle = 9°, slice thickness = 1.0 mm, FOV = 250 × 250 mm^2^ and a matrix of 256 × 256 for an effective spatial resolution of 0.97 × 0.97 × 1.0 mm^3^.

#### Quality control for MPRAGEs

Quality of structural scans for inclusion in a Voxel-Based Morphometry (VBM) analysis was determined through visual inspection conducted by three independent raters blind to participant group status (PS vs NS). Structural scans were scored on a 0 - 5 point scale for degree of artifacts that may affect the tissue classification algorithm (i.e., gross quality problems including phase wrap, ringing, and ghosting), where 0 indicates a scan free from visible artifact and 5 represents a scan of the poorest quality due to prominence of the relevant artifact. For each of the 3 raters, a weighted summary score was calculated. Summary rating scores were found to have good to excellent agreement between raters (intraclass correlation coefficient=.914, 95% confidence interval=.834-.950 [SPSS 24 based on mean rating (k=3), absolute agreement, 2-way mixed-effects model])^65^. The average weighted score of the 3 raters was assigned to each of the structural scans. Scans with the highest scores, indicating extremely poor quality, were flagged and after review by two of the authors (VLD and JVM) were confirmed to be unsuitable for VBM analysis.

#### MRI data preprocessing and analyses

Preprocessing for VBM was performed in SPM 8 (http://www.fil.ion.ucl.ac.uk/spm/software/spm8/). The *New Segment* option with default parameters identified different tissue types, generating native space tissue segmentation and DARTEL imported gray and white matter tissue maps. The DARTEL toolbox was used to create a study-specific whole-brain template (i.e., semi-optimized), and this template was registered to MNI space (affine transform) and modulated by the determinant of the Jacobian transform to preserve tissue volume and create images corresponding to gray matter volume (GMV). The GMV images were smoothed using a 12mm FWHM Gaussian kernel to reduce noise, compensate for spatial noise introduced during normalization, and improve validity of parametric statistics used^66^. To account for inter-individual differences in brain size, measures of GMV were normalized by total intracranial volume (ICV).

### Statistical analysis

#### Inclusion Criteria for Analyses

Given the potential impact of preterm birth^67^, low birth weight^68–70^, and exposure to alcohol and other substances *in utero*^71,72^ on fetal and postnatal brain development and behavior, analyses were restricted to adolescents whose imaging data met quality control criteria and were born at term (defined as >37 weeks) with normal birth weight (≥2500 grams, or 88.18 oz.) and no maternal report of tobacco, alcohol, regular prescription medication, or illicit drug use during pregnancy with the enrolled adolescent. Further, adolescent participants who reported substance use at baseline were excluded from analysis, given the effects of early substance initiation on adolescent neurodevelopment^73,74^. Finally, participants were excluded for poor image quality.

A total of 128 mothers completed the questionnaire. Based on the inclusion criteria detailed above, 45 participants were excluded from analyses. For details of participants remaining at each stage of data filtering, see Table S1, available online. Of the remaining 83 participants with complete pre-/perinatal history, 55 (66%) mothers reported having experienced no stressful event during pregnancy, and 28 (34%) reported having experienced at least one stressful life event during pregnancy (Table 2). Given the large number of respondents reporting no exposure to stressful events, groups were dichotomized into those whose mothers had not experienced stress during pregnancy (No-Stress group, *NS*; n = 55) and those who had experienced at least one event (Prenatal Stress group, *PS*; n = 28).

Seven mothers (NS = 4; PS = 3) reported presence of obstetric complications during pregnancy with the enrolled adolescent (Table 1). The mean birth weight for these 7 participants was 7.8 pounds (range: 6.9-8.8 pounds). Given that complications were reported in both groups, and infants were all full-term and born with normal birth weight^75^, these adolescents were included in analyses. Finally, 5 mothers (NS = 2; PS = 3) reported having been diagnosed with PPD after birth of the enrolled adolescent. As no additional details concerning PPD diagnosis were collected and the number of diagnosed mothers was small and distributed across both groups, we included these participants in analyses.

**Table 1.** Adolescent and maternal/perinatal characteristics

|  | **All Participants**<br>N = 83 | **Prenatal Stress (PS)**<br>N = 28 | **No Stress (NS)**<br>N = 55 | **p** |
| --- | --- | --- | --- | --- |
| *Adolescent characteristics* |  |  |  |  |
| Age at scan, Mean (SD) | 12.7 (.7) | 12.5 (.6) | 12.8(.7) | .061 |
| Sex |  |  |  | .051 |
| Males, n (%) | 38 (46%) | 17 (61%) | 21 (38%) |  |
| Females, n (%) | 45 (54%) | 11(39%) | 34 (62%) |  |
| Pubertal development Mean (SD) |  |  |  |  |
| Girls and boys | 2.27 (.72) | 2.32 (.73) | 2.25 (.72) | .693 |
| *Girls* | 2.56 (.69) | 2.93 (.64) | 2.44 (.68) | .042* |
| *Boys* | 1.97 (.54) | 1.93 (.49) | 2.0 (.59) | .69 |
| BMI z-score (SD) | .40 (.96) | .36 (1.1) | 42 (.86) | .762 |
| *Percentile (SD)<sup>a</sup>* | 61.3 (27.5) | 59.2(31.8) | 62.4(25.3) | .859 |
| Race (%) |  |  |  | .739 |
| *Caucasian* | 57% | 53.5% | 58.18% |  |
| *African American* | 30% | 32% | 29.1% |  |
| *Hispanic/Latino* | 6% | 3.5% | 7.27% |  |
| *Other* | 7% | 11% | 5.45% |  |
| Intelligence (K-BIT), Mean (SD) | 108.6 (15.7) | 107.1 (16.9) | 109.4 (15.1) | .529 |
| *Maternal/perinatal characteristics* |  |  |  |  |
| Maternal Age at delivery, mean (SD) | 30.9 (6.1) | 31.5 (7.5) | 30.6 (5.3) | .532 |
| Elapsed time, birth and maternal survey completion Mean (SD) | 13.3 (1.2) | 13.1 (1.1) | 13.4 (1.3) | .287 |
| SES index z-score (SD)<br>N = 80<br>*Parental education (years, mean (SD))*<br>*Household income (median)* | 16.0 (2.7)<br>\$100,000 - \$149,999 | n = 27<br>-.237 (1.06)<br>16.48 (2.7)<br>\$50,000 - \$74,999 | n = 53<br>.120 (.955)<br>16.48 (2.6)<br>\$100,000 - \$149,999 | .131 |
| Birth weight (ounces) Mean (SD)<br>N=79 | 122.8 (14.2) | n = 27<br>124.5 (15.1) | n = 52<br>121.9 (13.8) | .455 |
| Birth length, (inches) Mean (SD)<br>N = 58 | 19.7 (3.9) | n = 23<br>19.0 (6.1) | n = 35<br>20.2 (.9) | .261 |
| Obstetric anomaly (*n*)<br>*Description* | 7<br>*Gestational diabetes (n=4)*<br>*Bleeding during 1<sup>st</sup> trimester (n=1)*<br>*Breach/emergency C-section (n=1)*<br>*Unspecified (n=1)* | 3<br>*Gestational diabetes (n=2)*<br>*Breach/emergency C-section (n=1)* | 4<br>*Gestational Diabetes (n=2)*<br>*Bleeding during 1<sup>st</sup> trimester (n=1)*<br>*Unspecified (n=1)* | -- |
| PPD diagnosis (*n*)<br>*PPD therapy* | 5<br>*Medication (n = 2)*<br>*Therapy (n = 3)* | 3<br>*Medication (Sertraline (SSRI))(n=1)*<br>*Therapy (n=2)* | 2<br>*Medication (unknown) (n=1)*<br>*Therapy (n=1)* | -- |

Notes: SES = socioeconomic status, BMI = body mass index, PPD = postpartum depression, SSRI = selective serotonin reuptake inhibitor

#### Analyses

Dependent variable distributions were examined prior to analyses to confirm the data were free from outliers and that the assumption of normality was satisfied. The distribution of each of the eight DUSI-R subscales was non-normal (Shapiro-Wilk < .001, df = 82, for all subscales). Because standard transformations failed to normalize distributions, non-parametric statistical test were used for DUSI-R variables. Data analysis was conducted using SPSS Version 24.0.

PS and NS groups were compared on adolescent, prenatal, and demographic characteristics using Independent Samples Student’s *t*-test, Mann-Whitney U-test, and Chi-square test of independence, as noted. Additionally, groups were compared on the 8 DUSI subscales using the Mann-Whitney U-test with Bonferroni adjusted alpha levels of p < .00625 per test (.05/8) to correct for multiple comparisons.

#### Imaging

Statistical modeling compared PS and NS groups. Given that the distribution of males and females approximated a significant difference (p = .051) (Table 1), and because cortical thinning trajectories show sex differences^36–38^, sex was included as a covariate of no interest in imaging analyses. Clusters were defined using an uncorrected threshold of p < 0.001, k = 10. Corrections for multiple comparisons were made using non-stationary cluster correction (p < 0.05)^76,77^.

## RESULTS

### Group characteristics

Groups were similar in demographic (SES, race, and sex) and adolescent characteristics (age at MRI, IQ, pubertal development score, BMI, total intracranial volume, total GMV) (Table 1). Critically, groups were similar in neonatal characteristics (birth weight and length), allowing the inference that groups were similar in both neonatal head circumference^78^ and brain size^79^. Further, mothers were similar in age at delivery, and groups were similar in elapsed duration between the child’s birth and pre-/perinatal survey data collection. By design, the PS and NS groups differed regarding maternal experience of stress during pregnancy. Table 2 summarizes the stressful events reported by PS group mothers. Within the PS group, 6 mothers reported having experienced more than one event, (5 reported 2 events, 1 reported 3) (see Table S2).

**Table 2.**
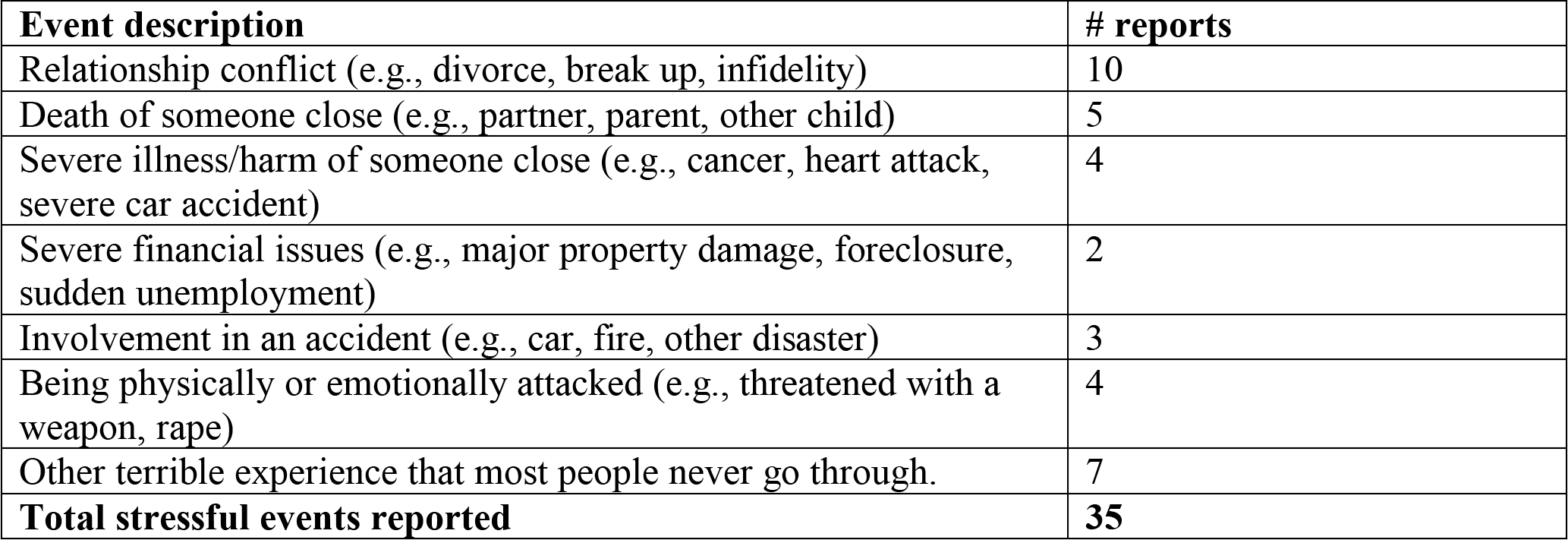
Summary of reported stressful events occurring during pregnancy in PS group.

| Event description | # reports |
| --- | --- |
| Relationship conflict (e.g., divorce, break up, infidelity) | 10 |
| Death of someone close (e.g., partner, parent, other child) | 5 |
| Severe illness/harm of someone close (e.g., cancer, heart attack, severe car accident) | 4 |
| Severe financial issues (e.g., major property damage, foreclosure, sudden unemployment) | 2 |
| Involvement in an accident (e.g., car, fire, other disaster) | 3 |
| Being physically or emotionally attacked (e.g., threatened with a weapon, rape) | 4 |
| Other terrible experience that most people never go through. | 7 |
| **Total stressful events reported** | **35** |

### Gray matter volume analysis

Whole brain non-stationary cluster corrected analysis revealed significantly greater GMV for the PS group compared to the NS group in bilateral parietal cortex (Table 3, Figure 1). Specifically, clusters in the right intraparietal sulcus (IPS: 40, −45, 55) (p_corr_ =.019), the left IPS (−32, −43, 55) (p_corr_ = .004), the left inferior parietal lobule (supramarginal gyrus) (IPL: −56, −23, 34) (p_corr_ = .035), and the left superior parietal lobule (SPL: −19, −44, 49) (p_corr_ = 0.048) showed greater volume for the contrast PS > NS (see Figures S2, S3). No significant results were found for the reverse contrast, NS > PS.

**Table 3.** Summary of results for contrast PS > NS, with sex as covariate of no interest. Coordinates are reported in MNI space. Results were thresholded at p<0.001, k =10 voxels. Cluster-level non-stationary corrected p-values are reported.

| Region | Cluster size | x | y | Z | Corrected p-value |
| --- | --- | --- | --- | --- | --- |
| R Intraparietal sulcus | 2093 | 40 | -45 | 55 | .019 |
| L Intraparietal sulcus | 1703 | -32 | -43 | 55 | .029 |
| L supramarginal gyrus | 1463 | -56 | -23 | 34 | .035 |
| L superior parietal lobule | 297 | -19 | -44 | 49 | .048 |

**Table 4.** Outcome measures

|  | All participants | Prenatal Stress (PS) | No Stress (NS) | p |
| --- | --- | --- | --- | --- |
| N | 83 | 28 | 55 |  |
| Mean Total ICV, Cubic Liters (SD) | 1.50 (.12) | 1.52 (.10) | 1.49 (.13) | .315 |
| Mean Total GMV, Cubic Liters (SD) | 0.72 (0.06) | 0.73 (0.05) | 0.72 (0.06) | .27 |
| **DUSI-R Subscales<sup>a</sup>** |  |  |  |  |
|  | Median (range) |  |  |  |
|  | 82 | 28 | 54 |  |
| Psychiatric symptoms | 20 (0-75) | 30 (0-75) | 15 (0-70) | .001<sup>*b</sup> |
| Family system | 7 (0-71) | 14 (0-71) | 7 (0-42) | .006<sup>*b</sup> |
| Behavior patterns | 15 (0-65) | 17 (0-65) | 10 (0-65) | .108 |
| Health status | 20 (0-50) | 20 (0-50) | 10 (0-50) | .033 |
| Leisure & Recreation | 16 (0-58) | 16 (0-58) | 12 (0-58) | .016 |
| School performance | 15 (0-60) | 17 (0-60) | 10 (0-45) | .010 |
| Social competence | 14 (0-64) | 21 (0-64) | 14 (0-57) | .046 |
| Peer relationships | 7 (0-42) | 14 (0-42) | 7 (0-42) | .033 |

Note: ICV = intracranial volume, GMV = gray matter volume, DUSI-R = Drug Use Survey Inventory, Revised
a. Mann-Whitney U-test used to examine group differences.
b. Significant at Bonferroni adjusted alpha level of $p < .00625$ .

**Figure 1.**
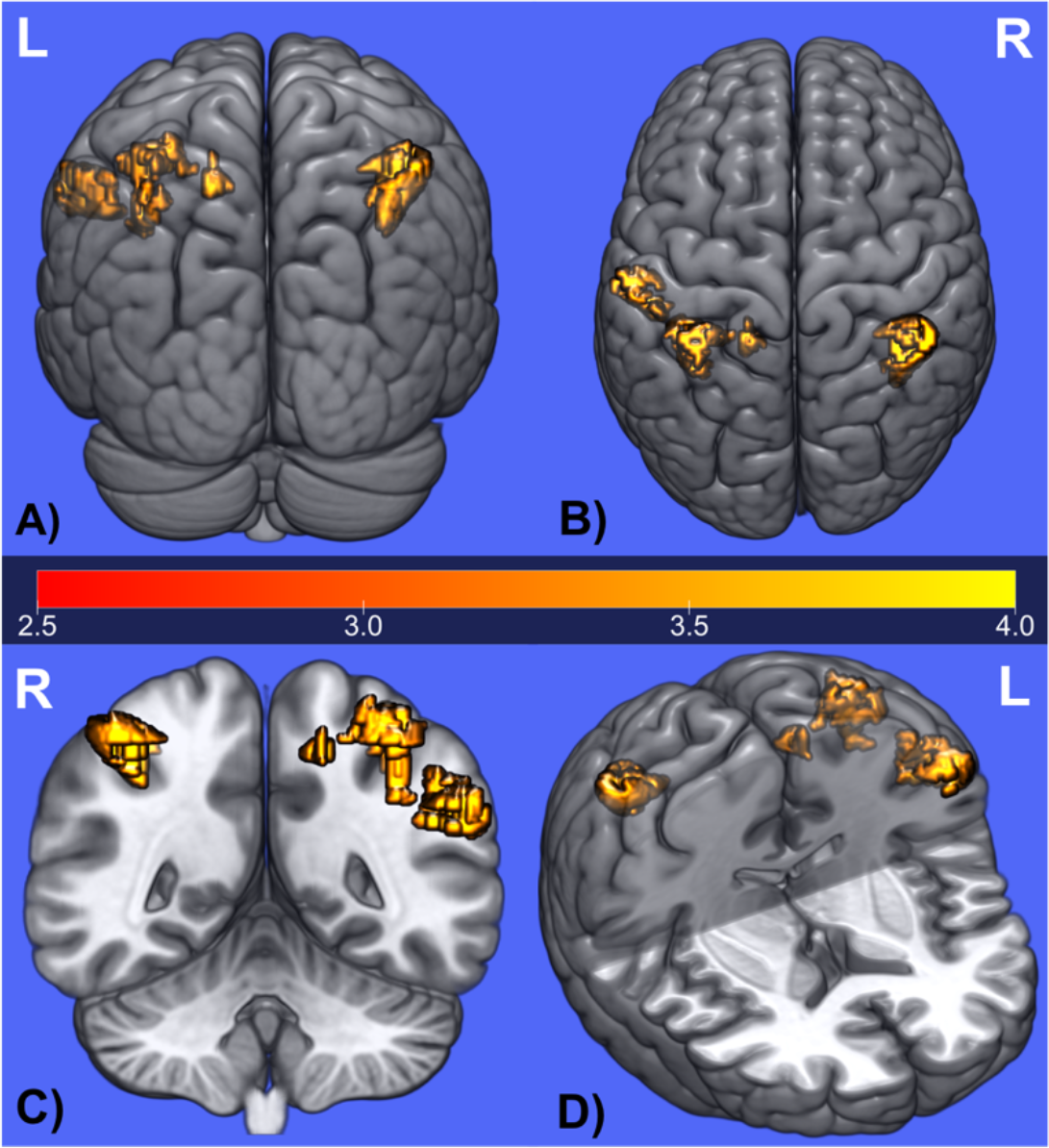
VBM analysis results for PS > NS using sex as a covariate of no interest; p < 0.05, non-stationary cluster corrected. Anatomical orientation showing posterior (A), superior (B), coronal (C) and cutaway (D) views of four PPC clusters (L = left, R = right).

### Risk measure

The two groups differed significantly on two of the eight DUSI-R subscales at the Bonferroni adjusted alpha-level of p < .00625 (Table 5, Figure 2). First, the PS group scored significantly higher on the DUSI psychiatric symptoms subscale (U(81)= 416.5,z=−3.339 p=.001), indicating a greater risk among the stress-exposed group for mental health symptomatology. Second, the PS group showed significantly higher scores for the family system subscale compared to the NS group (U(81) =486.5, z=−2.738, p=.006), indicating greater interfamilial dysfunction/conflict, less parental supervision, and/or less marital agility.

### Exploratory Analysis

To obtain a better understanding of the specific behavioral domains problematic for PS adolescents compared to their NS peers, we examined DUSI-R-derived subscales created specifically for use in screening for 4 psychiatric disorders in individuals under age 18^80^. The subscales, which probe features of ADHD, anxiety, depression and conduct disorder, cull questions from across the 10 original DUSI domains. These scales have been shown to be useful in identifying individuals who may require further evaluation for current psychiatric symptomatology, as well as those at risk for development of psychiatric disorder up to 8-10 years after their DUSI assessment^80^.

Scores were non-normally distributed and standard transformations were unsuccessful; we therefore compared the two groups using a one-tailed Mann-Whitney U-test, correcting for multiple comparisons using Bonferroni adjusted alpha levels of p < .0125 per test (.05/4). For all four DUSI-derived scales, PS adolescents showed higher scores, indictive of greater problems, and uncorrected values were significant for ADHD, anxiety, and depression. After corrections for multiple comparisons, only the ADHD scale remained significantly greater in PS adolescents, suggestive of greater risk for attentional dysfunction (U(81)= 506.5, z= −2.468, p=.007) (see Table S3).

## DISCUSSSION

To our knowledge, ours is the first study to examine the effects of exposure to maternal stress during gestation on brain morphology in typically-developing early adolescents. Compared to adolescents whose mothers report no exposure to a significant stressor during pregnancy, adolescents whose mothers experienced at least one stressful event showed increased GMV in PPC, and displayed greater risk for psychiatric symptoms and family system difficulties. Illuminating those effects of prenatal stress that may not manifest until adolescence is critical, given that latent deficits in structure may be revealed during this period of dramatic brain development,^39^ and moreover, altered developmental trajectories during this period may be associated with increased risk for negative outcomes^40,81^, including psychiatric symptomatology.

### Our central finding identified a relationship between maternal exposure to stress during pregnancy and GMV in early adolescence

The PS group exhibited increased GMV compared to the NS group in bilateral PPC: bilateral IPS, and left SPL and IPL. There were no regions for which NS adolescents showed increased GMV relative to the PS group. Changes in GMV from childhood through adulthood exhibit a heterochronic pattern, with different regions of cortex showing distinct developmental trajectories. Parietal gray matter increases during preadolescence, reaching maximum volume at 10.2 years for females and 11.8 years for males, and declining thereafter^36–38^. Given that groups were similar in chronological age (mean 12.5 years) and the potential influence of sex was controlled for, we would expect participants to be at similar stages in cortical development; that is, post-peak gray matter volume in PPC, with cortical pruning resulting in developmentally-appropriate cortical thinning^36–38,82,83^.

Our finding aligns with studies demonstrating a relationship between exposure to maternal distress during gestation and enduring effects on childhood brain morphology, including, importantly, in parietal cortex^31–33^. One notable difference between findings in the current study and findings of previous studies is the age of the children at the time of MRI scanning. While the present study examined early adolescents (11-14 years), previous studies examined children ages 6-9 or younger, a time in which normative development is characterized by increases in gray matter volume^36–38^. Thus, our results are in accord with previous research when considering this nonlinear developmental trajectory of the cortex. Our findings among early adolescents, taken together with the findings reported in the literature on early childhood, lead us to speculate that children exposed to the effects of stress *in utero* may show a temporal shift in peak gray matter volume, including a potentially altered trajectory of parietal cortical development, though this remains to be formally tested via longitudinal analyses.

We also found PS adolescents showed greater problematic behavior on the DUSI-R psychiatric symptoms subscale. Further, in exploratory analyses, the PS group had higher scores on the anxiety, depression, and ADHD subscales, though only the latter remained significant after corrections for multiple comparisons. These results are consistent with a robust literature showing an association between maternal stress during pregnancy and offspring cognitive and behavioral difficulties, including anxiety^21,84^, depression^23^, and schizophrenia^27,28^. Most relevant to the current study is evidence of an association between maternal stress exposure and impulsivity^22^ and ADHD^22,24,84,85^. Population cohort studies have demonstrated associations between maternal stressful life events such as those measured here and increased risk for ADHD in offspring. Boys born to mothers who experienced bereavement during pregnancy are at significantly increased risk for ADHD^86^. Similarly, maternal-rated ADHD symptoms in 2-year-olds is associated with exposure to stressful life events during pregnancy^87^.

Results from studies examining the relationship of ADHD and ADHD features to cortical development are consistent with the proposition advanced here that increased GMV in attentional processing regions is suggestive of a developmental lag in cortical development among PS adolescents. Youth with ADHD have been shown to exhibit the same region-specific pattern of brain development as TD youth, but that pattern is temporally delayed^82,88,89^; further, TD youth with ADHD features demonstrate a similar pattern shift, and increased ADHD features are associated with a more pronounced temporal shift. Among adults with childhood-diagnosed ADHD, only those who no longer meet ADHD diagnostic criteria exhibit normalization of cortical thickness in right parietal cortex, a key attentional region^88^.

Consistent with its diverse structural and functional connectivity patterns, the PPC is a component of multiple brain networks, and PPC subregions are hubs in both dorsal and ventral fronto-parietal attention networks^90–92^. The connections between PPC regions with other structures is key, as input provided by PPC could have an impact on downstream structures, particularly frontal regions. Interestingly, many of the regions that previous studies report as having what may be interpreted as delayed development ^21,31,33^ are connected to PPC, including frontal cortex, and meta-analytic research has shown hypoactivation in the frontoparietal network in children with ADHD^93^. Although the PS group in the current study scored higher in ADHD-type features, they also demonstrated increased depressive and anxious symptoms relative to the NS group. Thus, problems with ADHD-type features may more broadly reflect difficulties in executive function, which could be suggestive of a non-specific risk for psychiatric features, given that deficits in executive function are attested in many disorders (e.g., depression, anxiety, and schizophrenia).

The findings reported here must be interpreted within the context of limitations of the current study. First, along with increased psychiatric symptoms, the PS group showed significantly higher scores on the family system subscale of the DUSI-R, a 14-item subscale querying parental involvement/supervision, parent-imposed boundaries/rules, occurrence of frequent or intense arguments between the adolescent and his/her parents or others in the home, as well as questions concerning regular or problematic substance use by or arrest of an immediate family member. We are unable to disentangle the contribution of prenatal maternal stressful life events from that of current dynamics within the child’s family since many of the factors in our measure of prenatal stress could foreshadow difficulties in postnatal familial dynamics. Second, the questionnaire-based measure of prenatal stress has limitations. Retrospective assessment of traumatic events may include biases^94^. However, measures such as the one used here have been shown to be accurate and reliable^95–97^, and the two groups were similar regarding elapsed time between the adolescents’ birth and questionnaire completion. Also, the questionnaire did not include an assessment of the mothers’ perception of how stressful the event was. The subjective distress associated with an event rather than the objective event itself has been associated with level of trauma among first-responders in a natural disaster^98^. However, among pregnant women it has been shown that objective events (even in the absence of maternal report of subjective distress during the event) are associated with sequelae among offspring^18,99^. Additionally, the questionnaire did not inquire about the timing of the stressful event during pregnancy, though it is unclear whether gestational stress experienced earlier^100,101^ or later^31^ is more impactful, and it has been suggested timing of the stressor is not a critical factor in offspring outcomes^102^.

Strengths of the present analysis include the rich characterization of the two, relatively large groups of typically developing adolescents analyzed, including descriptors related to prenatal, postnatal, maternal and adolescent variables. Existing studies asking similar research questions have been correlational, have had smaller total sample sizes, and did not possess the rich characterization of participants presented in the current study^31,32^. Importantly, the chronological age range of the adolescents studied here was narrow and thus more readily allows for consideration of group differences during this period of dynamic alterations in gray matter. Additionally, factors known to be related to GMV, including age, IQ^103^, pubertal development,^36^ BMI^104^, and sex^36,37^, were either controlled for or were similar between groups in the current study.

Given the normative development of parietal cortex, increased GMV in the PS group is potentially suggestive of a developmental lag in cortical maturation compared to their similarly aged NS peers. Our results add to the growing literature demonstrating the enduring effects of maternal stress exposure on brain morphology and the possibility of increased risk for psychiatric symptoms. It is plausible that delayed development in PPC may impact structures that rely on PPC input, creating a feedforward effect, resulting in differential input to other developing structures, especially the prefrontal cortex, which shows a protracted course of development^83,105^. Adolescence is a key period in brain and psychosocial development, during which the risk for emergence of neuropsychiatric disorders is heightened^40^ and realized^106,107^. The groups studied here differed neither for proxies for estimated brain size at birth, nor for ICV and total GMV at the time of MRI scans, suggesting that despite overall similarities in brain structure both at birth and in early adolescence, region-specific differences in brain development resulting from *in utero* stress exposure are evident in adolescence, a key neurodevelopmental period. It is therefore critical to understand the factors that contribute to differences in the course of brain development during adolescence, and how these differences potentially increase risk for the development of psychiatric symptoms. Understanding the enduring impact of maternal prenatal stress exposure on adolescent brain and behavior provides an opportunity to target those youth at increased risk for problems and identify factors that can help them attain positive functional outcomes. Furthermore, these results highlight the importance of providing pregnant women with sufficient social support to help mitigate the effects of external stressors.

