## Supplemental Materials for "Altered cortical development and psychiatric symptom risk in adolescents exposed to maternal stress *in utero*"

*Table S1. Participants remaining after each stage of data filtering based on inclusionary/exclusionary criteria*

| | *N* |
|---|---|
| 1. Enrolled in study | 135 |
| 2. Completed prenatal questionnaire | 128 |
| 3. Excluded at prenatal questionnaire/demographics check: | **25**[*] |
| Preterm birth | 5 |
| Low birth weight | 1 |
| Exposure to substance *in utero* | 20[*] |
| Alcohol consumption | 6 |
| Marijuana use | 2 |
| Nicotine use (smoking) | 6 |
| Prescription medication regularly used | 8 |
| Adolescent baseline substance use | 3 |
| 4. Passed to MRI quality check | 103 |
| 5. Excluded due to image quality | **20** |
| **Final sample** | **83** |

**Note:** *Number of participants excluded for specific reasons may not sum to totals because of participants who failed multiple inclusionary/exclusionary criteria.*

*Table S2. Summary of stressful events reported by mothers in the PS group who experienced more than one event.*

| **Event description** | **# reports** |
|---|---|
| Relationship conflict (e.g., divorce, break up, infidelity) | 4 |
| Death of someone close (e.g., partner, parent, other child) | 3 |
| Severe illness/harm of someone close (e.g., cancer, heart attack, severe car accident) | 1 |
| Severe financial issues (e.g., major property damage, foreclosure, sudden unemployment) | 0 |
| Involvement in an accident (e.g., car, fire, other disaster) | 1 |
| Being physically or emotionally attacked (e.g., threatened with a weapon, rape) | 2 |
| Other terrible experience that most people never go through. | 2 |
| **Total stressful events reported** | **13** |

*Table S3. DUSI-R-derived psychiatric scales*

| | **All participants** *(N = 82)* | **Prenatal Stress (PS)** *(n=28)* | **No Stress (NS)** *(n=54)* | ***p*** |
|---|---|---|---|---|
| | | **Median (range)**[a] | | |
| ADHD | 20 (0-80 ) | 35 (0-70 ) | 20 (0-80) | .007[b] |
| Anxiety | 20 (0-66) | 33 (0-60) | 20 (0-66) | .0145 |
| Depression | 16 (0-83) | 22 (0-55) | 11 (0-83) | .0255 |

| | | | | |
|---|---|---|---|---|
| Conduct Disorder | 5<br>(0-47) | 5<br>(0-42 ) | 2<br>(0-47) | .1105 |

*a. Mann-Whitney U-test used to examine group differences.*
*b. Significant at Bonferroni-corrected p < .0125*

*Figure S1. DUSI-R problem density scores for PS and NS groups. Higher scores are indicative of greater problems within that domain. PS group shows higher scores in all domains, and after Bonferroni correction for multiple comparisons the PS group significantly differs from NS group on psychiatric symptoms and family system subscales (p < .00625).*

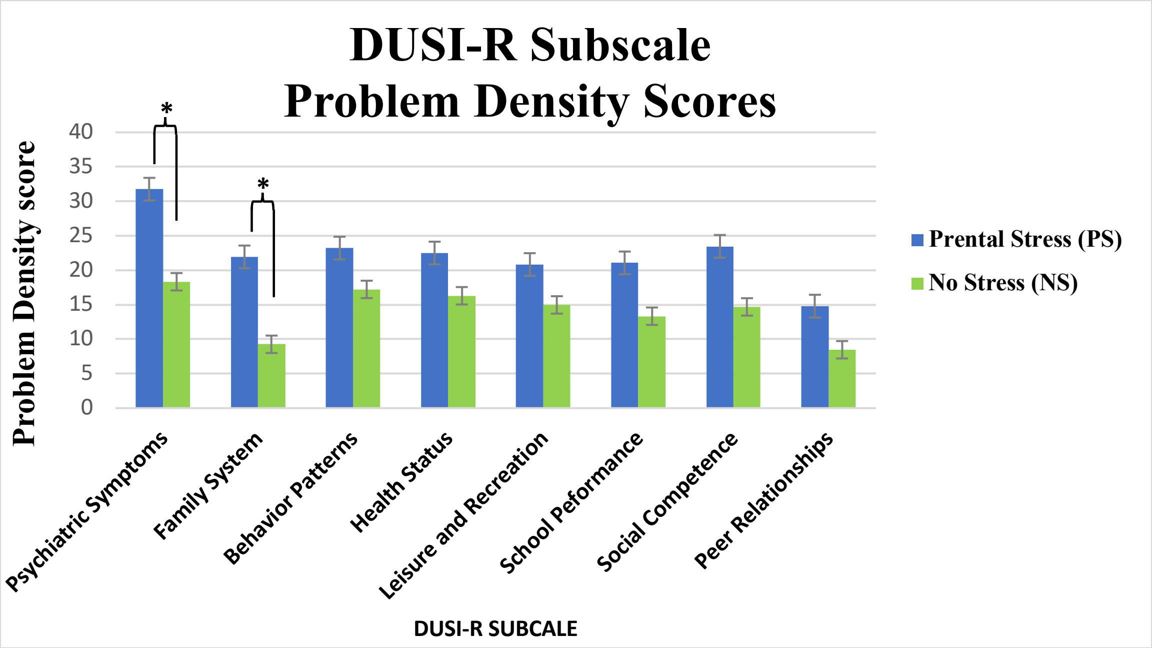

*Figure S2. GMV in four posterior parietal regions for which PS group showed greater GMV relative to NS group. A) Left SPL, B) Left SMG, C) Left IPS, D) Right IPS; SPL = superior parietal lobule, SMG = supramarginal gyrus, IPS = intraparietal sulcus.*

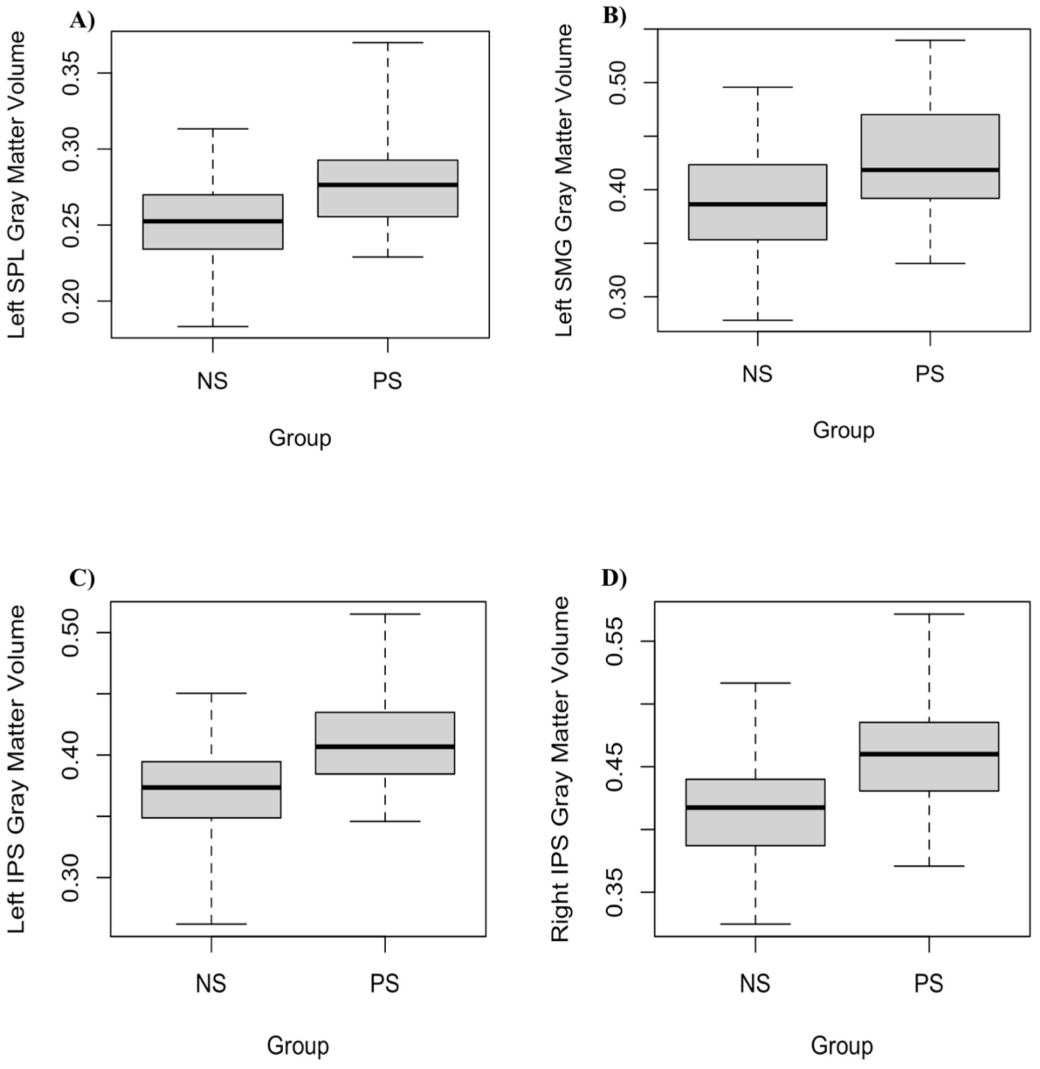

*Figure S3. Total mean GMV in the four posterior parietal cortex (PPC) clusters for which PS adolescents showed increased GMV compared to NS adolescents.*

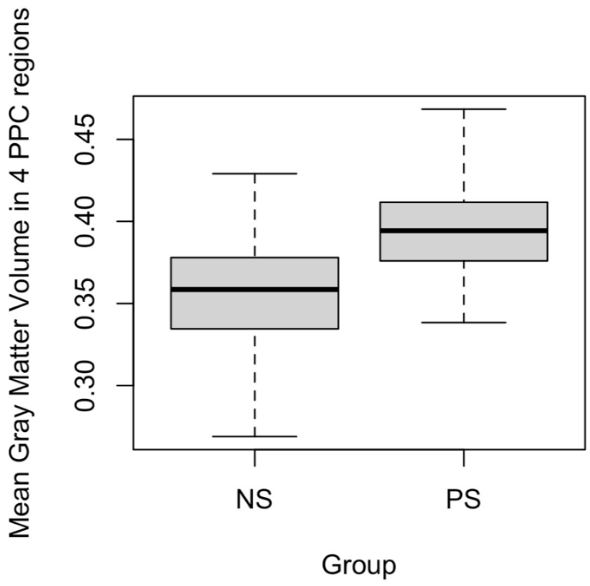
